# Interaction Between the Prefrontal and Visual Cortices Supports Subjective Fear

**DOI:** 10.1101/2023.10.23.562918

**Authors:** Vincent Taschereau-Dumouchel, Marjorie Côté, Shawn Manuel, Darius Valevicius, Cody A. Cushing, Aurelio Cortese, Mitsuo Kawato, Hakwan Lau

## Abstract

It has been reported that threatening and non-threatening visual stimuli can be distinguished based on the multi-voxel patterns of hemodynamic activity in the human ventral visual stream. Do these findings mean that there may be evolutionarily hardwired mechanisms within early perception, for the fast and automatic detection of threat, and maybe even for the generation of the subjective experience of fear? In this human neuroimaging study, we presented participants (Fear group: N=30; No Fear group: N = 30) with 2700 images of animals that could trigger subjective fear or not as a function of individual’s idiosyncratic “fear profiles” (i.e., fear ratings of animals reported by a given participant). We provide evidence that the ventral visual stream may represent affectively neutral visual features that are statistically associated with fear ratings of participants, without representing the subjective experience of fear itself. More specifically, we show that patterns of hemodynamic activity predictive of a specific “fear profile” can be observed in the ventral visual stream whether a participant reports being afraid of the stimuli or not. Further, we found that the multivariate information synchronization between ventral visual areas and prefrontal regions distinguished participants who reported being subjectively afraid of the stimuli from those who did not. Together, these findings support the view that the subjective experience of fear may depend on the relevant visual information triggering implicit metacognitive mechanisms in the prefrontal cortex.

## Introduction

Recently, using multivoxel pattern analysis of magnetic resonance imaging (fMRI) data, it was found that one can decode or classify from the patterns of activity in the human visual cortex between threatening and non-threatening visual stimuli seen by the subjects [1,2]. This has led to the intriguing claim that there may be emotional schemas embedded within the human ventral visual system [2].

Taken further, perhaps one provocative interpretation could be that representations of fear itself could be found within the ventral visual stream, reflecting evolutionarily hard-wired mechanisms for the purpose of automatic detection of threat [3,4]. However, an alternative interpretation could also be that, threatening stimuli (i.e., stimuli that some individuals interpret as threatening and likely to generate fear [4–7]), may, statistically, share certain visual features. For instance, some commonly feared animals and insects are likely to share certain shapes and surface texture, such as scales or shells. Accordingly, what the early visual processes represent may not be a prioritized processing of fear-associated visual features or even the representation of subjective fear *per se*, but rather, only objective visual properties that generally or statistically predict fear.

To arbitrate between these two different interpretations, we can find stimuli that are only reported to be subjectively fearful to some human participants, but not others, such as commonly feared animals. That way, we can experimentally dissociate between objective visual stimuli and subjective fear as indicated by self-report by individual subjects. Importantly, using such an approach, we can test if the “fear profile” (i.e., subjective fear ratings of different animal categories reported by a specific participant) of participants reporting subjective fear (“Fear” group) can also be decoded based on fMRI patterns of activity in participants reporting no subjective fear (“No fear” group). If such decoding turns out to be equally accurate in both groups, this may support the hypothesis that the ventral visual stream only represents neutral visual features typically associated with fear, but not subjective experience of fear *per se*.

In this study, participants were presented with a set of 2,700 images of 30 commonly feared animals and provided fear ratings of each of the animal categories. Two groups of participants were created based on their fear profiles: those reporting at least two high-fear ratings (“fear” group; N=30) and those reporting no high-fear rating of any animals (“No fear” group; N = 30). We investigated differences between these groups by conducting brain decoding analyses in the ventral visual stream as a whole and within 4 different subregions: the occipital, fusiform, inferior temporal and middle temporal gyri (based on the Brainnetome atlas [8]). In the “fear” group, we trained within-subject machine learning decoders to predict the fear profile of participants based on the brain activity generated by individual images. In the “No fear” group, we predicted the fear profiles of participants in the “Fear” group within the same brain regions and compared decoding performance with those observed in the “Fear” group (see Fig. 1C). By comparing decoding performance between both groups, we aim to determine if the ventral visual cortex represents affectively neutral visual features or the subjective experience itself. As such, if decoding performances are not statistically different between the groups, this would indicate that participants in the “Fear” group are not representing any information that goes beyond what is otherwise available during the processing of the same visual stimuli in other participants.

**Figure 1.**
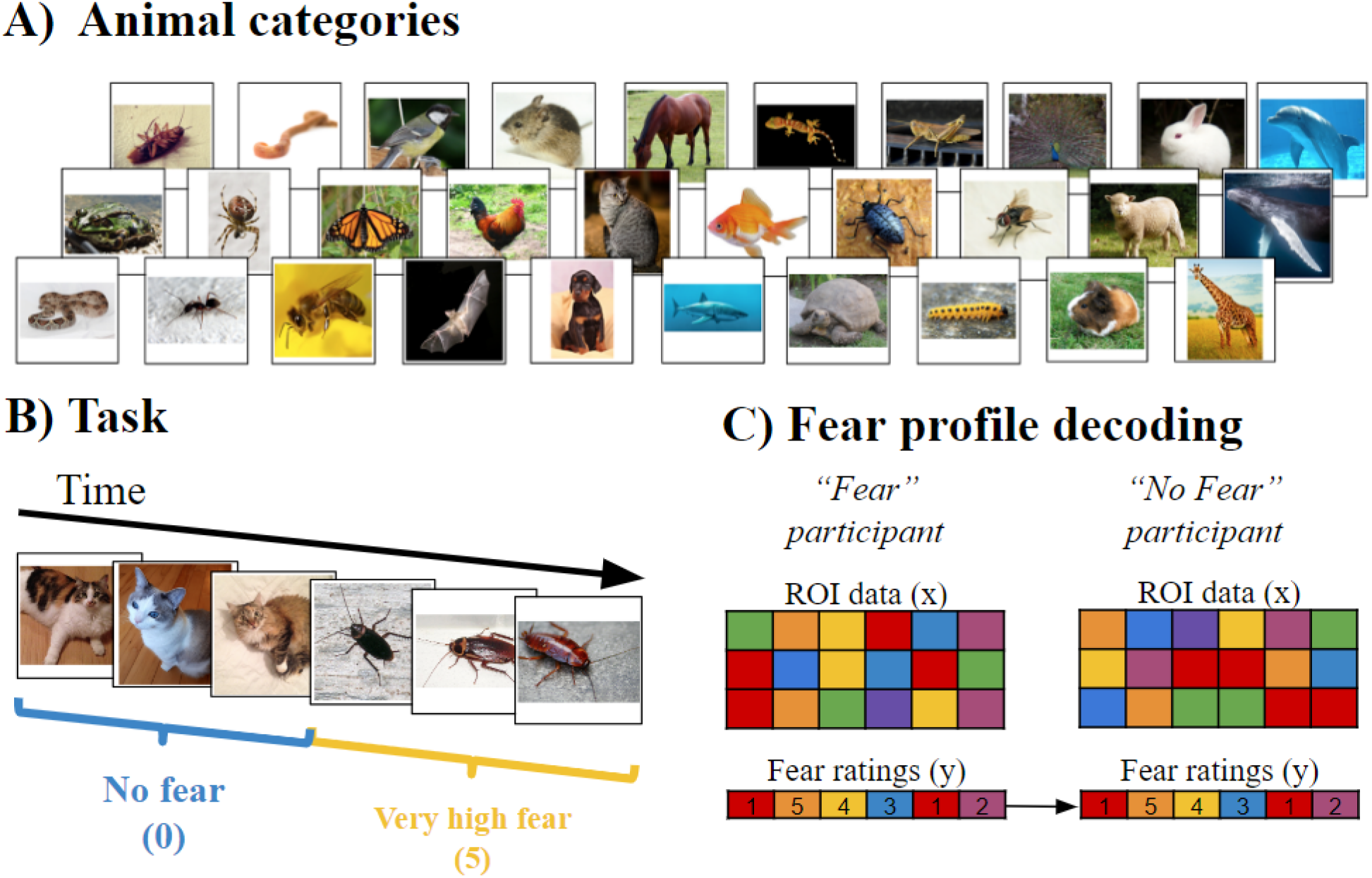
(A) Animal categories included in the fMRI experiment (see text for a complete list). (B) Participants were presented with a series of 3600 images of animals and human-made objects, each lasting 0.98 s. They were asked to pay attention to the image category and report any category change (e.g. from ‘cat’ to ‘cockroach’ as shown in the figure) with a button press. (C) Participants reporting high fear of some animals in the dataset, presented a unique “fear profile”. Those profiles were decoded using (1) the participants’ brain data (“Fear” participant) or (2) the brain data of other participants that were also presented with the same images (“No Fear” participants). The decoding of the fear profile of each participant in the “Fear” group was compared to the mean decoding of that specific fear profile in the 30 participants in the “No Fear” group.”

To further investigate how fear profiles could be predicted from affectively neutral visual features, we conducted a series of analyses using artificial neural networks. Many artificial neural networks have now reached human level recognition on a great variety of visual stimuli [9,10]. The information represented within the different layers of those networks has often been mapped to the visual system and is commonly used as a model of the information processing in the ventral visual stream [11,12]. Interestingly, we can determine the pattern of activity generated in pre-trained and fixed networks by a new set of images. Such patterns of activity are often termed “latent-space representations” or “embeddings” and they can be analyzed using machine learning approaches. If we use pre-trained networks that are not particularly trained to recognize fear profiles, but are instead trained to recognize images in general, the embeddings of the images are likely to reflect visual properties of the images. As such, if we can predict the fear profiles of participants in the “Fear” group solely from the embeddings of the images in our database, this would be a further indication that above chance decoding results can be obtained solely based on neutral visual features. To do so, we decoded the fear profiles of the “Fear” group from the embeddings of the 2,700 images in 2 pre-trained artificial neural networks: a transformer-based vision model (Contrastive Language-Image Pretraining; CLIP) [10] and a deep convolutional neural network (Visual Geometry Group 19; VGG19) [9]. VGG19 comprises 19 layers of varying dimensions (see methods for more details) while the latent-space of the CLIP model includes 512 dimensions. We used the information contained in these embeddings to decode fear profiles. Above chance decoding of fear profiles in those artificial neural networks could further support the notion that fear profiles can be decoded from neutral visual features alone.

Furthermore, we investigated if any difference between the “Fear” and “No fear” groups arise in the communication between the ventral visual stream and other brain areas. In fact, multiple theoretical perspectives suggest that subjective experience may be linked to a broad system of brain regions lying outside the ventral visual stream [4,5,7,13–16]. Such regions include the amygdala [17,18], the hippocampus, the anterior cingulate cortex, the insula and various subregions of the prefrontal cortex [19]. We reasoned that subjective fear experience may actually be associated with the communication of the affectively neutral information represented in the ventral visual stream with such brain regions. For this purpose, we segmented the amygdala, hippocampus, parahippocampus, insula and the main regions of the prefrontal cortex using the Brainnetome atlas [8] and conducted information synchronization analyses [20] between the ventral visual cortex and each of these brain regions. Information synchronization analysis can be thought of as a multivariate connectivity metric. It can be used to determine how decoded information in a ventral visual area is synchronized with another brain region, thus indicating an association between the represented information in the two brain regions. By comparing information synchronization between the two groups, we hope to determine where in the information processing hierarchy the subjective experience arises.

## Methods

### Participants

Thirty participants (fourteen females, mean age 23.3 ± 4.35 years) were recruited to take part in an fMRI experiment at the *ATR (Advanced Telecommunications Research)-Computational Neuroscience Laboratories* in Japan. Participants were recruited if they presented self-reported “high” or “very high” fear of at least one animal in our database using a 6-point Likert scale. Amongst this group, 3 participants were diagnosed with a specific animal phobias using the Structured Clinical Interview for DSM-IV. Thirty additional participants were also selected from a larger cohort (N = 53) of participants that underwent the same fMRI experiment (see *Study design*). These participants were selected to act as a control group for the purpose of the current study and were included if they presented no “high” or “very high” fear of any animals included in the dataset (3 females, mean age 23.1 ± 2.87 years). For both groups, inclusion criteria were: (a) aged between 18 and 45; (b) no psychotropic medications; (c) no contraindication to magnetic resonance imaging. The inclusion criteria were specified on the recruitment advertisements and verified through screening forms and an additional assessment on the first day of the study. The study was approved by the ATR Research Ethics Board and the participants provided informed written consent.

### MRI parameters

Participants had their brain hemodynamic signals measured and recorded in two 3T MRI scanners (Prisma Siemens and Verio Siemens) with a 32-channels head coil at the ATR Brain Activation Imaging Center. During the experiments, we obtained 33 contiguous slices (TR = 2000 ms, TE =30 ms, voxel size = 3 × 3 × 3.5 mm 3, field-of-view = 192 x 192 mm, matrix size = 64 x 64, slice thickness = 3.5 mm, 0 mm slice gap, flip angle = 80 deg) oriented parallel to the AC-PC plane, which covered the entire brain. We also obtained T1-weighted MR images (MP-RAGE; 256 slices, TR = 2250 ms, TE = 3.06 ms, 5 voxel size = 1 × 1 × 1 mm 3, field-of-view= 256 x 256 mm, matrix size = 256 x 256, slice thickness = 1 mm, 0 mm slice gap, TI = 900 ms, flip angle = 9 deg.).

#### Stimuli presentation in the fMRI scanner

Visual stimuli were projected on a translucent screen using an LCD projector (DLAG150CL,Victor). The projected image spanned 20 × 15 deg in visual angle (800 × 600 resolution) and had a refresh rate of 60 Hz. The experiment presentation was conducted using the PsychoPy2 software (v1.83) [21] and images covered 13.33 degrees of visual angles during the procedure.

#### Study design

Participants were presented with 3,600 pictures of animals and objects grouped in mini-blocks of 2, 3, 4 or 6 images of the same basic category. Trials were organized into six runs of 600 trials interleaved with short breaks. Each image was presented for 0.98 seconds. To make sure that participants paid attention to image categories, they were asked to report any change in category (e.g. from one kind of animal to another) by pressing a button using their right hand. The sequence of image presentation was pseudo-randomized and fixed across participants. In order to allow high-pass filtering of the fMRI data, chunks within each category were organized so that their period was always shorter than 120 seconds.

We included 90 images of each of the animal and object categories. The 30 animal categories included reptiles (snake and gecko), amphibians (frog and turtle), insects (cockroach, beetle, ant, spider, grasshopper, caterpillar, bee, butterfly, and fly), birds (robin, peacock, and chicken), annelids (earthworm), mammals (mouse, guinea pig, bat, dog, sheep, cat, rabbit, horse, and giraffe) and aquatic animals (shark, whale, common fish, and dolphin). The human-made objects included: airplane, car, bicycle, scissor, hammer, key, guitar, cellphone, umbrella, and chair. The data from the human-made objects were not analyzed in the current project as we focused on animal fear. The 3600 images were collected from various sources on the Internet, including: the Creative Common initiative (https://creativecommons.org), Pixabay (images marked for commercial use and modifications; http://pixabay.com), Flickr (images allowing commercial use and modifications; http://www.flickr.com), and Shutterstock (http://shutterstock.com). The images were selected if they presented a full frontal view of the object or animal and if no other objects were clearly identifiable in the background. Images were cropped so that they would frame the object. The final images were 533 × 533 pixels and covered 13.33 degrees of visual angles during the procedure. The average contrast and luminance of images were not different between categories (see supplementary material of [22]).

## Data Analysis

### Data pre-processing

MRI results included in this manuscript come from preprocessing performed using fMRIPrep 1.5.9 ([23]; RRID:SCR_016216), which is based on Nipype 1.4.2 ([24,25]; RRID:SCR_002502).

### Functional data preprocessing

For each of the 6 BOLD runs per subject, the following preprocessing was performed: First, a reference volume and its skull-stripped version were generated using a custom methodology of fMRIPrep. Susceptibility distortion correction (SDC) was omitted. The BOLD reference was then co-registered to the T1w reference using bbregister (FreeSurfer) which implements boundary-based registration [26]. Co-registration was configured with six degrees of freedom. Head-motion parameters with respect to the BOLD reference (transformation matrices, and six corresponding rotation and translation parameters) are estimated before any spatiotemporal filtering using mcflirt (FSL 5.0.9, [27]). BOLD runs were slice-time corrected using 3dTshift from AFNI 20160207 ( [28], RRID:SCR_005927). The BOLD time-series (including slice-timing correction when applied) were resampled onto their original, native space by applying the transforms to correct for head-motion. These resampled BOLD time-series will be referred to as preprocessed BOLD in original space, or just preprocessed BOLD. The BOLD time-series were resampled into standard space, generating a preprocessed BOLD run in [‘MNI152NLin2009cAsym’] space. A reference volume and its skull-stripped version were generated using a custom methodology of fMRIPrep. Gridded (volumetric) resamplings were performed using antsApplyTransforms (ANTs), configured with Lanczos interpolation to minimize the smoothing effects of other kernels [29]. Many internal operations of fMRIPrep use Nilearn 0.6.1 (Abraham et al. 2014, RRID:SCR_001362), mostly within the functional processing workflow. For more details of the pipeline, see the section corresponding to workflows in fMRIPrep’s documentation.

NiLearn [30] was used to detrend, remove motion confounds (24 parameters: 3 rotations, 3 translations, their time derivatives, power 2, and derivative power 2) and standardize data. Single-trial estimates were then obtained using the least-square separate approach [31,32] implemented using functions from pyMVPA [33,34]. This method allows to iteratively fit a general linear model to estimate the brain response to each image. Each general linear model includes one parameter modeling the current trial and two parameters modeling all other trials in the design.

### Decoding fear profiles in the ventral visual stream

We used single-trial estimates of brain activity to predict the reported level of fear within-participants (0 = “No fear” to 5 = “Very high fear”). Across all animal categories, participants in the Fear group presented higher fear ratings (M = 1.55; SD = 0.64) than participants in the “No Fear” group (M = 0.52; SD = 0.28). Since the distribution of fear ratings tended to be skewed (i.e. many participants reported a disproportionate number of categories eliciting “No Fear”), we randomly under-sampled the “No Fear” level to match the mean number of trials in other fear levels. This was achieved to prevent introducing a bias in the models and to ensure a proper proportion of Fear trials (of all fear levels) in the training dataset. After under-sampling, the mean number of trials was 1857.2 ± 612 trials.

Decoding was achieved using a 6-fold cross-validation, as a function of experimental runs, using least absolute shrinkage and selection operator (LASSO) regression (e.g., [35] as implemented in Scikit-Learn [36]. To prevent overfitting, the alpha parameter was first tuned, ROI per ROI, in a nested cross-validation fashion in 5 participants and averaged. Those values were used to conduct further analyses. We used the coefficient of determination (R2) as a measure of performance and the Fisher-transformed correlation coefficient between the predicted and real values are also presented in Fig. S1. Decoding was conducted within the entire ventral visual stream and separately within 4 subregions: occipital cortex, fusiform gyrus, inferior temporal gyrus and middle temporal gyrus. The regions of interest were determined as a function of the Brainnetome Atlas annotation [8]. Masks of the 4 ventral visual regions are illustrated in Fig. 2.

**Figure 2.**
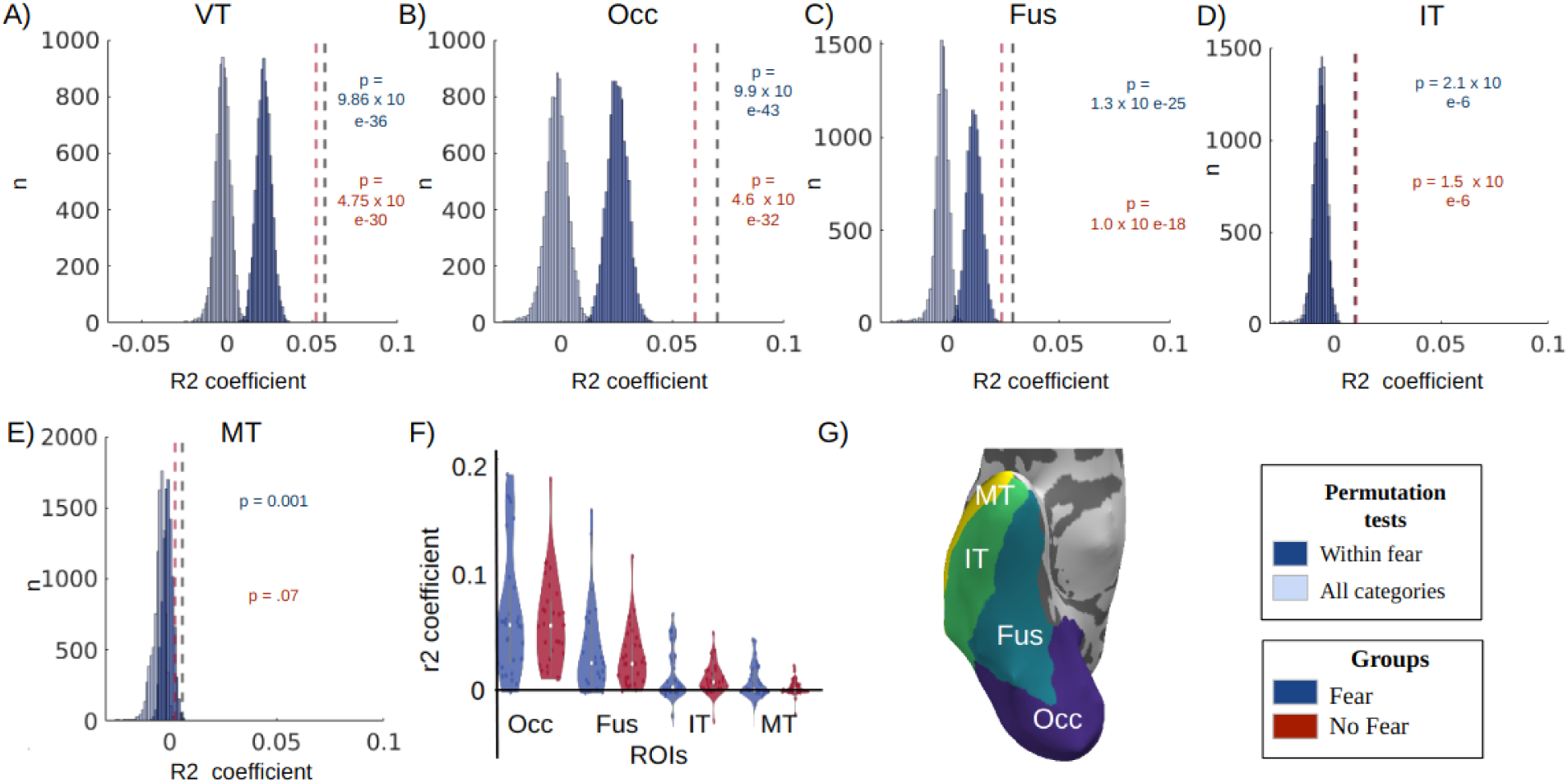
Prediction of the fear profiles in participants with (“Fear group”) and without (“No fear” group) subjective fear of the animals. A-E) Generally, the fine-grained spatial patterns of hemodynamic activity in the entire ventral visual stream (VT) and within all four subregions (occipital cortex, Occ; fusiform gyrus, Fus; inferotemporal cortex, IT, and middle temporal cortex, MT) can distinguish, better than chance, between images of threatening and non-threatening animal categories (p-values are computed with respect to the permutation of all categories; see main text for statistical information). This is shown by comparing mean decoding performance, within each group, against two random distributions of group means obtained by conducting decoding of (1) randomly permuted category labels (light blue) and (2) randomly permuted category labels within high-and low-fear ratings independently (dark blue; see text for more details). F) No group differences were observed, indicating that above chance decoding is obtained regardless of whether the human participants in questions reported being subjectively afraid of the typically threatening animal categories. This dissociation between subjective fear and stimulus threat was possible because some ‘threatening’ animals (e.g. cockroaches) were only fearful to some but not all participants. Violin shapes represent density and dots individual participants (Fear group) or group mean (No Fear group). Central dot represents the mean and error bars’ edges the 1st and 3rd quantiles. G) Regions of interest based on the Brainnetome atlas and displayed using pySurfer (https://github.com/nipy/PySurfer/).

Two permutation tests were used in order to determine above chance performances. In the first permutation test, we randomly shuffled (1000 times) the fear profiles of participants in the Fear group at the Category level and decoded their brain activity in each ROIs. This permutation test was conducted to determine if just any combination of categories could be decoded with the same accuracy. Similarly, we conducted a second permutation test, this time permuting (1000 times) the fear ratings within the high-fear categories (>=4 on the Likert scale) and low-fear categories (<4 on the Likert scale) independently. This second permutation test was conducted to determine if the fearful and non-fearful categories could be permuted without affecting decoding performances. In both permutation tests, group performances were determined by randomly sampling (10,000 times) the permuted values for each participant and computing new group averages. This created random distributions of group averages (Shown in Fig. 2) that were used to determine the statistical significance of the real group means. Those tests were corrected for multiple comparisons using the Bonferroni correction (4 ROIs).

Decoding performances in the “Fear” group were also compared to the decoding of the same fear profile in the “No fear” group. This was achieved in order to determine if the same decoding performance could be obtained in participants reporting no subjective fear of the presented animals. In order to do so, we predicted the fear ratings of each participant in the “Fear” group from the brain activity of each of the 30 participants in the “No fear” group (i.e., 30 × 30 decoders). Paired-sample t-tests were used to compare the mean predictions of a given fear profile in the “No Fear” group to the prediction of the corresponding participant in the “Fear” group. The Bonferroni correction was used to control for multiple comparisons (4 ROIs) and the Bayesian paired-sample t-tests were used to determine the likelihood of rejecting the null hypothesis. Those results are presented in Fig. 2f.

We conducted a control analysis to estimate the effect of sex imbalance in the “No Fear” group as only a small proportion of the group was female (3 females). In order to estimate the effect of sex imbalance we created a sub-group in the “No Fear” group that was balanced with respect to sex (3 men and 3 women) by randomly sub-sampling the participants in those groups. By comparing the mean of this subgroup to the Fear group using paired-sample t-tests we hoped to estimate if sex imbalance could explain our results. The Bonferroni correction was used to control for multiple comparisons (4 ROIs). Those results are presented in Fig. S2.

We also conducted another control analysis to rule out the effect of fear profiles similarity. Even if participants in the “No fear” group do not report high-fear ratings of animals in our dataset, it is still possible that above chance decoding could be achieved if their fear profiles are correlated with the fear profiles of participants in the “Fear” group. To rule out this possibility, we subsampled the “No Fear” participants to be included for a specific fear profile comparison. More specifically, for the comparison to each fear profile in the “Fear” group, we excluded the participants in the “No fear” group whose fear profile presented a significant correlation with the fear profile of the target participant. This way, each mean only included participants presenting no correlation in their fear profile. We conducted paired-sample t-tests to compare those means with the prediction of the corresponding participant in the “Fear” group. The Bonferroni correction was used to control for multiple comparisons (4 ROIs). Those results are presented in Fig. S2.

### Decoding fear profiles from image embeddings in deep neural networks

We also aimed to determine if deep neural networks trained to recognize images could be used to predict the fear profiles of participants. We used two different networks with different architectures: a deep convolutional neural network (Visual Geometry Group 19; VGG19) [9] and a transformer-based vision model (Contrastive Language-Image Pretraining; CLIP) [10]. For both networks, we used pre-trained versions of the models, propagated our images in these neural networks and trained decoders to predict each of the 30 fear profiles of participants in the Fear group based on the activity generated in each layer of the networks (see details below).

We used the “imagenet-vgg-verydeep-19” version of VGG19 from the MatConvNet website (https://www.vlfeat.org/matconvnet/) trained on the ILSVRC-2012 dataset that included 1,000 image categories of various animals and human-made objects. It includes 19 layers: 16 convolutional and 3 fully connected layers: Conv1 (Conv1_1 and Conv1_2; 3211264 units), Conv2 (Conv2_1 and Conv2_2; 1605632 units), Conv3 (Conv3_1 and Conv3_2; 802816 units), Conv4 (Conv4_1,Conv4_2, Conv4_3, and Conv4_4; 401408 units), Conv5 (Conv5_1, Conv5_2, Conv5_3 and Conv5_4; 100352 units), fc6 (4096 units), fc7 (4096 units) and fc8 (1000 units) (For more details on the network, see [9]). The MatConvNet toolbox for Matlab [37] was used in order to extract the image embeddings.

We used the “openai/clip-vit-base-patch32” version of CLIP available on Hugging Face (https://huggingface.co/openai/clip-vit-base-patch32). Briefly, CLIP is designed to learn visual concepts and their associated textual descriptions by training on a large corpus of images and corresponding textual descriptions from data found on the internet. The model is trained in a contrastive manner that leverages both image and text embeddings to establish meaningful associations between images and their corresponding textual descriptions. It is based on the Vision Transformer (ViT) architecture, a popular model for image classification tasks. The "clip-vit-base-patch32" variant utilizes a patch size of 32×32 pixels for processing images. Here, we extracted the latent-space embedding of each image, after projection to the latent space (512 dimensions).

The embeddings of our 2700 images in the two networks (i.e., in each layer of VGG19 and in the latent space of CLIP) were used to train machine learning decoders to predict the fear profile (i.e., fear ratings of a given participant to each of the 30 animal categories) of the 30 participants in the “Fear” group. For the image embeddings of CLIP, a LASSO regression was implemented in a 6-fold cross-validation framework (as a function of experimental runs) with an alpha parameter determined using nested cross-validation in 5 participants. Performances were determined using the coefficient of determination between the predicted and real fear rating values. Significance was determined using the same 2 permutation tests as described in the section “Decoding fear profiles in the ventral visual stream”.

A similar approach was used to determine the prediction capacity of each layer within VGG19. However, since some layers included a great number of units (e.g., 3211264 for Conv1_1 and Conv1_2), we elected to use partial-least square regression as implemented in Scikit-Learn [36] in order to first decrease the dimensionality of the data. Performances were also determined using the Fisher-transformed correlation coefficient and significance was determined using one-sample t-tests corrected for multiple comparisons using the Bonferroni correction. Those results are presented in Fig. S3.

### Image synthesis based on the embedding decoders

In pyTorch, we carried out a procedure to generate latent-space embeddings corresponding to high outputs of specific fear profile decoders. The optimization process included 300 iterations in order to update an initial zero vector in latent space as a function of the loss function computed between a high fear value and the predicted value by the latent-space decoder. As a result, the zero vector was iteratively updated using the backpropagation of this error. The resultant latent-space embeddings were then reconstructed visually using the Stable UnCLIP pipeline available on Hugging face (https://huggingface.co/docs/diffusers/api/pipelines/stable_unclip). This approach allows to leverage Stable Diffusion 2 (https://huggingface.co/docs/diffusers/api/pipelines/stable_diffusion/ <u>stable_diffusion_2</u>) in order to generate visual images conditioned on the CLIP vision embeddings [38]. This procedure was used in order to synthesize the visual features leading to high outputs of the latent-space fear profile decoders.

### Information synchronization with other brain regions

We used information synchronization analysis [20] to determine between-group differences in the communication of the ventral visual regions with other brain regions. Essentially, this analysis determines if decoded information in a seed region (i.e., predicted fear ratings in the fusiform region) is synchronized (i.e., correlated) with decoded information in another brain region. As a result, this analysis can indicate the synchronization between two brain regions if their decoded information is correlated. This was achieved using(LASSO regression (e.g., [35] as implemented in Scikit-Learn [36]. To prevent overfitting, the alpha parameter was first tuned, ROI per ROI, in a nested cross-validation fashion in 5 participants and averaged. Those values were used in the remaining analyses. Fisher-transformed correlation coefficients were used as a measure of synchronization. When the algorithm did not converge, a constant value was output by the decoders. In those situations, synchronization scores were not computed for these specific participants. This resulted in a number of missing participants (up to 8) for the estimation of some synchronization analyses. For completeness, we also report synchronization results without any tuning of the alpha parameter (fixed value of 0.1) and without missing participants in Fig. S4.

Target” regions were selected based on the segmentation of the Brainnetome atlas [8]. We used the segmentation of the prefrontal cortex to establish the dorsolateral prefrontal cortex (middle frontal gyrus), the ventrolateral prefrontal cortex (inferior frontal gyrus), orbitofrontal and ventromedial cortex (orbital gyrus). The medial prefrontal cortex and anterior cingulate cortex was defined using the anterior part of the cingulate gyrus and the anteromedial part of the superior frontal gyrus. The insula (insular gyrus), amygdala, para-hippocampal and hippocampal cortex also followed the segmentation of the Brainnetome atlas. Those regions were included for their alleged role in affective information processing [1]. We compared the mean synchronization results in the “No Fear” group to the corresponding participants in the “Fear” group using paired-sample t-tests. Significance was determined after correcting for multiple comparisons using the False discovery rate approach [39].

## Results

### Decoding fear profiles in the ventral visual stream

When compared to the random permutation of animal categories, the true fear profiles were predicted above chance in the ventral visual stream (p = 1.18 x 10e-3; t(29) = 5.822, p = 2.597 × 10e-6, Bonferroni corrected, Mean R2 = 0.057, STD = 0.054, Cohen’s d = 1.056) as well as in the 2 subregions namely the occipital cortex (p = 3.797 × 10e-43; t(29) = 7.09, p = 3.36 × 10e-7, Bonferroni corrected, Mean = 0.07, STD = 0.054, Cohen’s d = 1.300) and the fusiform gyrus (p = 1.3 × 10 e-25; t(29) = 4.33, p =.0002, Bonferroni corrected, Mean = 0.03, STD = 0.037, Cohen’s d = 0.81). The two other sub-regions, the inferior temporal gyrus (p = 2.1 × 10 e-6; t(29) = 1.68, p = 0.418, Bonferroni corrected, Mean = 0.0099, STD = 0.032, Cohen’s d = 0.30) and the middle temporal gyrus (p =.001; t(29) = 2.10, p = 0.045, Bonferroni corrected, Mean = 0.0058, STD = 0.015, Cohen’s d =0.387) presented mixed results.

Fear profiles were not predicted more accurately in participants reporting subjective fear of the animals compared with participants reporting no fear in any of the 4 regions (“Fear” group vs “No Fear” group; Occipital: t(29) = 1.353, p = 0.187, Bonferroni corrected; Fusiform: t(29) = 1.159, p = 0.256, Bonferroni corrected; Inferotemporal: t(29) =-0.069, p = 0.945, Bonferroni corrected; Middle temporal: t(29) = 1.571, p = 0.127, Bonferroni corrected). Bayesian paired t-test indicated no evidence to reject the null hypothesis in the Occipital cortex (BF_10_ = 0.443), Fusiform gyrus (BF_10_ = 0.358) and inferior temporal gyrus (BF_10_ = 0.195) and the middle temporal gyrus (BF_10_ = 0.584) [40–42].

Furthermore, no group effect can be found after including in the “No Fear” group only participants without correlation in their fear profiles with the targeted participant in the “Fear” group (Occipital: t(29) = 1.950, p = 0.244, Bonferroni corrected; Fusiform: t(29) = 1.188, p = 0.978, Bonferroni corrected; Inferotemporal: t(29) =-0.553, p = 1.0, Bonferroni corrected; Middle temporal: t(29) = 1.981, p = 0.228, Bonferroni corrected). Similarly, no group differences can be observed when groups are balanced with respect to sex (Occipital: t(29) = 1.461, p = 0.620, Bonferroni corrected; Fusiform: t(29) = 1.12, p = 1.0, Bonferroni corrected; Inferotemporal: t(29) =-0.002, p = 1.0, Bonferroni corrected; Middle temporal: t(29) = 1.937, p = 0.250, Bonferroni corrected).

### Predicting fear profiles from image embeddings in deep neural networks

The image embeddings in the latent space of the CLIP network could be used to predict, above chance, the 30 fear profiles of the participants in the “Fear” group (p = 1.13 × 10e-6, t(29) = 37.404; p = 4.2862 × 10e-26). The image embeddings in the different layers of VGG19 networks can also be used to predict, above chance, the 30 fear profiles of our participants (see Fig. S3). The t-values ranged between 5.6620 (fc2: t(29) = 5.6620, p = 2.04 × 10e-04; Bonferroni corrected) and 7.203 (conv5_2: t(29) = 7.203 6.10 × 10e-06; Bonferroni corrected) with the Conv5 layers presenting the highest coefficients (Mean = 0.3631 to 0.3846; STD = 0.249 to 0.263). Only FC3 did not present a significant prediction of the fear profiles (t(29) = 1.62; p = 0.120; Bonferroni corrected).

### Information synchronization with other brain regions

Information synchronization analyses were conducted using the 3 ventral visual stream ROIs as seed regions. Participants in the “Fear” group showed a greater information synchronization between the occipital gyrus and two regions in the prefrontal cortex, namely the ventromedial prefrontal cortex (t(27) = 2.89, p = 0.0075; significant after FDR; BF_10_ = 5.82) and the medial prefrontal cortex / anterior cingulate cortex (t(27) = 3.095, p = 0.0044; significant after FDR; BF_10_ = 9.07). Similarly, participants in the “Fear” group showed a greater information synchronization transmission between the fusiform gyrus and the insula (t(21) = 3.14, p = 0.0049; significant after FDR; BF_10_ = 8.73) and the dorsolateral prefrontal cortex (t(21) = 2.92, p = 0.0081; significant after FDR; BF_10_ = 5.64). Lastly, participants in the “Fear” group also showed a greater information synchronization between the inferior temporal gyrus and the ventromedial prefrontal cortex (t(21) = 2.92, p = 0.0081; significant after FDR; BF_10_ = 5.65).

## Discussion

In summary, as in some previous studies [1,2], here we found that “fear profiles” can be predicted from patterns of hemodynamic activity generated by threatening and non-threatening stimuli in the human visual and visual association cortices. However, this was the case regardless of whether the human subjects reported to be subjectively afraid of the visual stimuli in question. Further, even if the fear profiles presented idiosyncratic variability between participants, we still found that they could be predicted from the activity patterns within artificial neural networks that were not trained to identify threat or fear *per se* (but rather, just to identify different objects). Based on the information captured by the decoders of the artificial neural network CLIP, we generated synthetic stimuli to illustrate the visual information distinguishing threatening and non-threatening stimuli. Interestingly, these generated stimuli also do not seem to look subjectively threatening. Together this seems to support the hypothesis that the early visual representations do not actually encode fear, but rather, just visual features that are statistically common in stimuli that can be interpreted as threatening by some individuals.

In contrast, the main positive finding is that subjective fear was reflected by information synchronization between different prefrontal regions and ventral visual areas, specifically the occipital and fusiform gyrus, and to a lesser extent, the inferotemporal area (IT). This is to say, participants who reported to be subjectively afraid of the relevant animals showed heightened information synchronization in these pathways as they watched the threatening stimuli. This finding may add some credence to the view that subjective experiences require implicit metacognitive processes that depend on the prefrontal cortex [5–7,13].

Notably, we did not observe this difference in information synchronization from ventral visual areas to the amygdala. This area has traditionally been thought to be important for fear processing [17,18]. However, much of the evidence behind that idea came from studies of animal models, most notably in rodents [43–46]. In such studies, fear is only indirectly inferred based on physiology or behavior. In a recent study in humans, we have also found that physiological arousal (i.e., skin conductance response) in reaction to viewing threatening stimuli can in fact be predicted by patterns of hemodynamic activity in the amygdala [1]. However, self-reports of subjective fear were better predicted by patterns of hemodynamic activity in prefrontal areas [1]. These results are also in line with other recent fMRI studies indicating that the subjective experience of fear [47] and threat anticipation [48] are best predicted when brain decoders are not restricted to the amygdala alone.

We also did not observe a significant difference in information synchronization between ventral visual areas and the ventrolateral prefrontal cortex. This prefrontal region receives input from the ventral visual areas, especially IT. In a recent study, it was found that chemical inactivation of this prefrontal region in monkeys can impair object recognition, as it dampens feedback responses to IT [49]. However, this mechanism seems to concern objective identification in general, especially in ambiguous images, but not directly affective processing.

Together, these findings could perhaps be considered under Tulving’s distinction between anoetic, noetic, and autonoetic conscious processing [50–52]. The information flow from ventral visual areas to amygdala may be considered anoetic (lacking knowledge), as it likely reflects physiological responses that aren’t specific, with respect to visual content. The information flow to the ventrolateral prefrontal cortex may be considered noetic (knowing), but it concerns the information about the visual objects rather than oneself. It is the interaction between the ventral visual stream and other prefrontal areas, including ventromedial prefrontal, medial prefrontal, and dorsolateral prefrontal cortices, that reflects autonoetic processes, i.e. processes about oneself [53]. It has been argued that fear as a conscious experience always requires self-related mechanisms [6,53]. While the current study provided information supporting this view of autonoetic consciousness, future studies will be needed to investigate the role of the amygdala and ventrolateral prefrontal cortex in anoetic and noetic consciousness respectively.

The current study has several important limitations. For example, the threatening visual stimuli are all animals. In real life, there are of course other kinds of threatening stimuli, such as weapons. It is possible that images of animals are processed by evolutionarily hardwired mechanisms, and therefore differently from other inanimate stimuli. It remains to be tested in future studies whether the current findings would generalize.

Also, we did not directly assess the nature of the decoded information in the ventral visual stream of both groups. As such, it is still possible that the visual system of participants in the “Fear” group simply do not represent those visual features in the same way. However, based on previous analyses, we do not expect this hypothesis to be likely [22]. Previously, we used a functional alignment method called hyperalignment, to determine if the brain activity in the ventral visual stream of a group of surrogate participants could be used to train animal decoders that could generalize to participants presenting diverse fear profiles and even to patients diagnosed with specific phobia (see [22], supplementary Fig. 6). Our results indicate that “hyperalignment decoders” performed with the same accuracy whether they were tested on participants with different fear profiles or on patients presenting specific phobia. As such, these results indicate that representations in the ventral visual stream are unlikely to vary considerably between the “Fear” and “No Fear” groups. However, future studies could expand on these results and determine if the same pattern of results could also be observed when decoding the fear profiles of participants instead of individual animal categories.

Also, our key positive findings depend on the analysis of information synchronization. This analytic method is not totally new, and variants of the approach have been employed in numerous previous studies [22,54–56]. It focuses on how information as captured by patterns of hemodynamic activity (rather than overall level) is reflected by patterns of activity in another region. In this sense, it is a slightly more advanced multivoxel variant of standard connectivity analysis. However, like standard connectivity analysis, it is a correlational method. For understanding causal interactions between brain areas, invasive interventional methods are more powerful and rigorous. Unfortunately, they are not easily employed in human studies. Future studies on animal models can address this issue better.

Another limitation of the current study is our reliance on a subjective criterion to determine group membership. As this study was not a clinical trial, there was not an obvious way to determine group membership (for instance, patients vs controls). As such, we aimed to remain mostly consistent with what was used in previous studies(Taschereau-Dumouchel et al., 2018, 2020). However, this limits the usefulness of the results as they cannot provide much information regarding mental health conditions such as specific animal phobias. Other studies will be needed to address these questions directly.

Finally, in assessing the subject fear level in response to the synthetic images generated by the artificial neural network models (Fig. 3), we did not conduct formal behavioral tests. We only visually inspected the images ourselves, and feel that such formal tests are not necessary, because the images barely resemble the actually threatening images. Also, this is not a main finding for the current study. However, we cannot preclude the existence of subtle arousal effects. We plan to address this limitation in a future study. If these synthetic stimuli are proven not to elicit an excessive level of fear or discomfort, even in patients with phobia of the relevant animals, one interesting possibility may be to test if these synthetic stimuli can be used for the purpose of exposure therapy-without the patients having to directly encounter the unpleasantness of seeing the actual images of the phobic objects.

**Figure 3.**
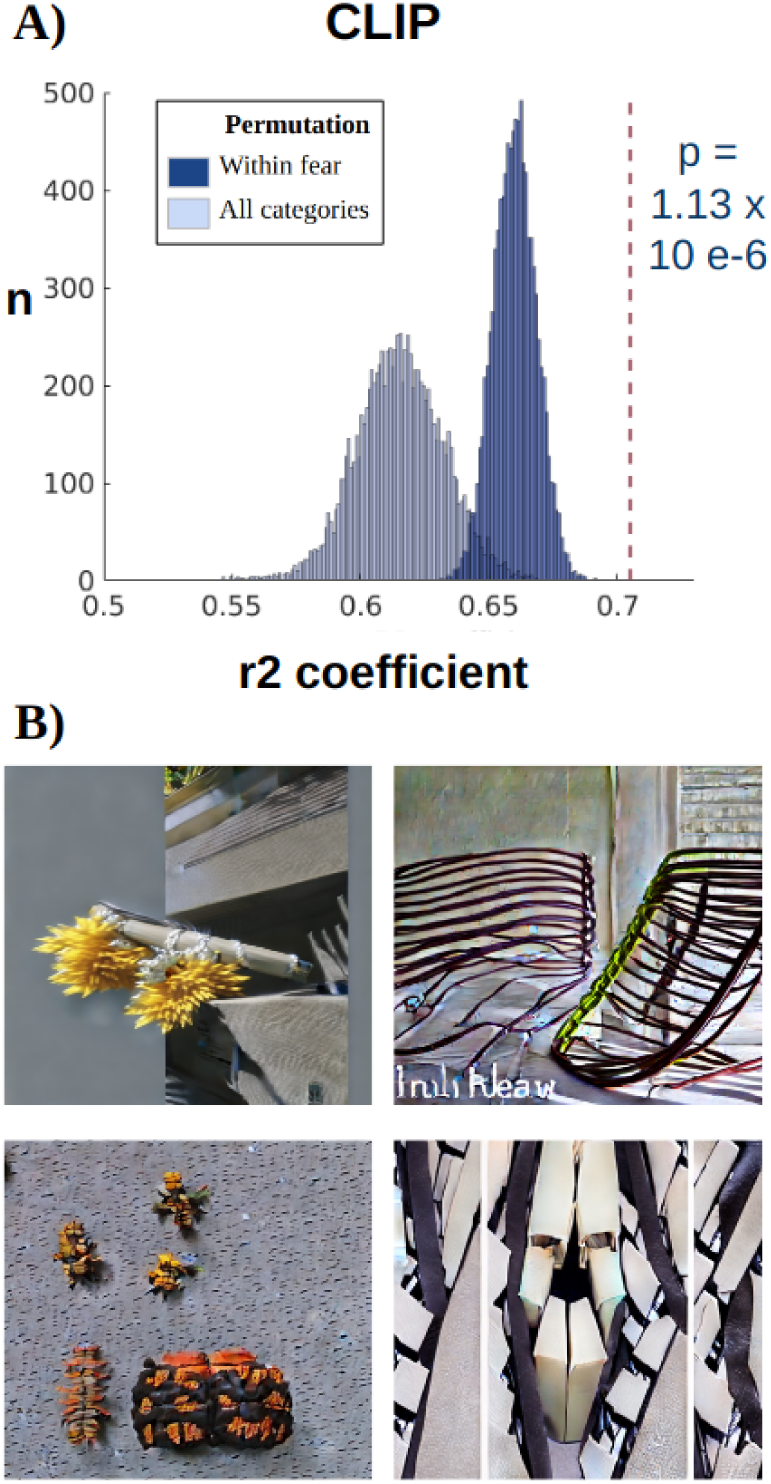
(A) Fear profiles of participants can be predicted from the activity generated by the 2700 images in theartificial (‘deep’) neural networks: CLIP(the vision ‘transformer’). By fear profile we mean the different self-reported subjective fear scores over all the animal categories, for an individual participant. Based on the pattern of activity in “latent space”within the artificial neural network over many stimuli, we tried to predict these fear profiles for each participant. The r2 coefficient is a measure of how well activity from the ‘latent-space’ of the network (see main text for more details), can accurately predict the fear profile over different animal categories. These results indicate that CLIP can perform far better than chance (see main text for statistics). (B) Synthetic images generated using the decoders of fear profiles of 4 participants (based on the CLIP embeddings). To understand the nature of the relevant representations within these networks that allowed the above results, we used an optimization procedure and StableUnCLIP to generate synthetic images that represent the ‘prototypical’ content for some fear profiles of participants. As one can see, these synthetic images do not necessarily resemble animals but include visual features of some of the most feared animals in the participants’ profile (from left to right, bee, worm, caterpillar and spider). Based on our own subjective inspection, the synthetic images do not necessarily appear to be fear-inducing.

**Figure 4.**
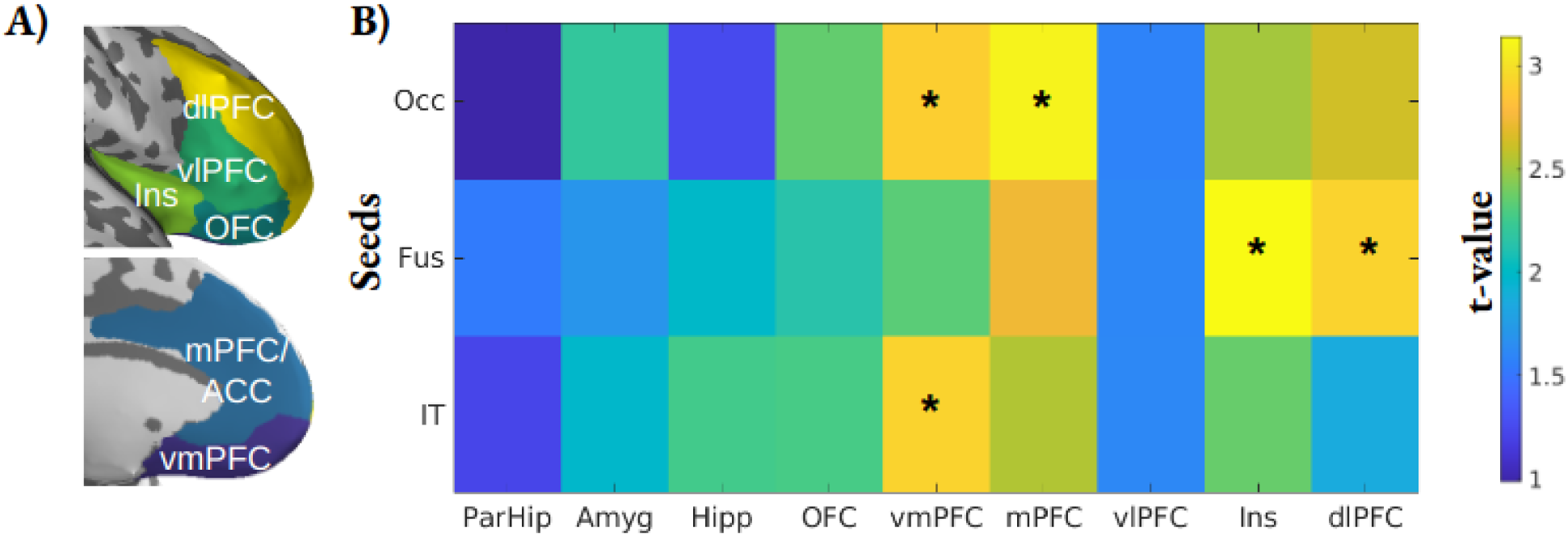
Difference in information synchronization between ventral visual regions and other brain areas, between participants with and without subjective fear of ‘threatening’ stimuli. Color coded represent the t-value of the between group difference in a measure of information synchronization. The measure essentially captures how the multivoxel pattern in a seed region (Occ, Fus, IT; same label as used in Figure 2), with respect to the degree to which it can distinguish between threatening vs non-threatening stimuli, can be predicted by the multivoxel pattern in another “target” region (para-hippocampal area, ParHip; amygdala, Amyg; hippocampus, Hipp; orbitofrontal cortex, OFC; ventromedial prefrontal cortex, vmPFC; medial prefrontal cortex, mPFC; ventrolateral prefrontal cortex, vlPFC; insula, Ins; dorsolateral prefrontal cortex, dlPFC). Specifically, what is plotted is not the absolute value of information synchronization, but rather the difference in these values between participants who reported to be afraid of the relevant threatening stimuli, and participants who reported not to feel so. Marked in asterisks (*) are pathways that are significantly different between the two groups of participants, after Bonferroni correction (see main text for statistical details). In other words, these information synchronization pathways distinguished between different levels of self-reported subjective fear (across participants), while the physical stimuli (including both images of typically threatening and non-threatening animal categories) were held constant. The image of the ROIs was generated based on the Brainnetome atlas using pySurfer (https://github.com/nipy/PySurfer/).

## Acknowledgements and funding statement

V.T-D. was supported in part by the *Fond de recherche du Québec-Santé* and the *Fondation de l’Institut universitaire en santé mentale de Montréal*. A.C. and M.K. are partially supported by the Japan Science and Technology agency ERATO Ikegaya brain-AI fusion (grant number JPJMER1801), by JSPS KAKENHI (grant number JP22H05156), and by the Agency for Technology, Labour and Innovation (grant number JP004596).

## Data and code availability

Data and codes to recreate the statistical analyses can be found here: https://osf.io/5xtgc/?view_only=b7f4fbc85ddc4fbf8fc7074412b1e3ff

## Notes

### Competing Interest Statement

The authors have declared no competing interest.

### Summary of Updates

Inclusion of permutation tests to the main analyses; use of synchronization analyses instead of transmission analyses.

