## Supplementary material for "Interaction Between the Prefrontal and Visual Cortices Supports Subjective Fear"

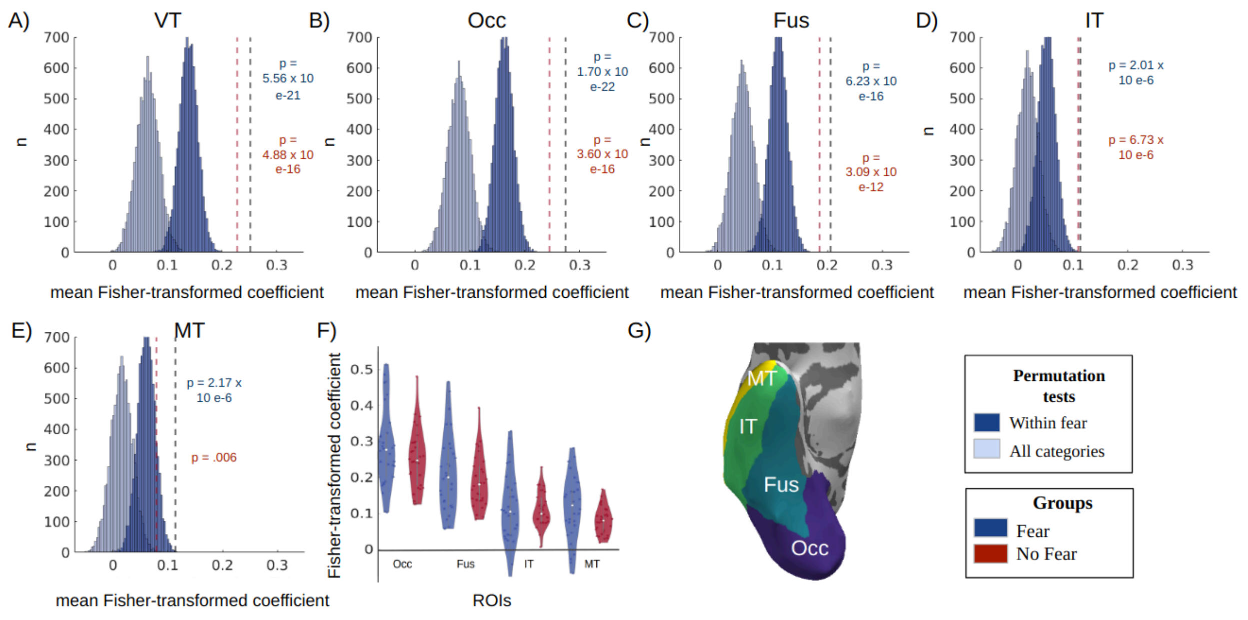


**Figure S1. Prediction of the fear profiles in participants with ("Fear group") and without ("No fear" group) subjective fear of the animals.** This figure recreates the same analyses as presented in Fig. 2 but with Fisher-transformed correlation coefficients instead.

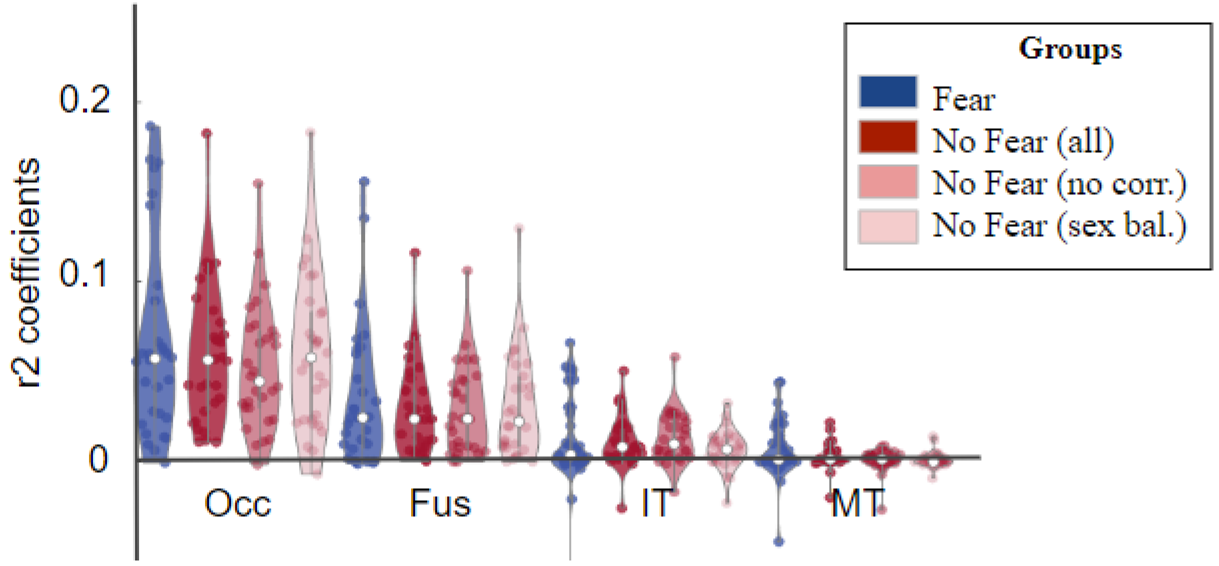


**Figure S2.** No group effect can be found when only participants without correlation in their fear profiles are included in the "No fear" group (see main text for statistical detail). Similarly, no group differences can be observed when groups are balanced with respect to sex.

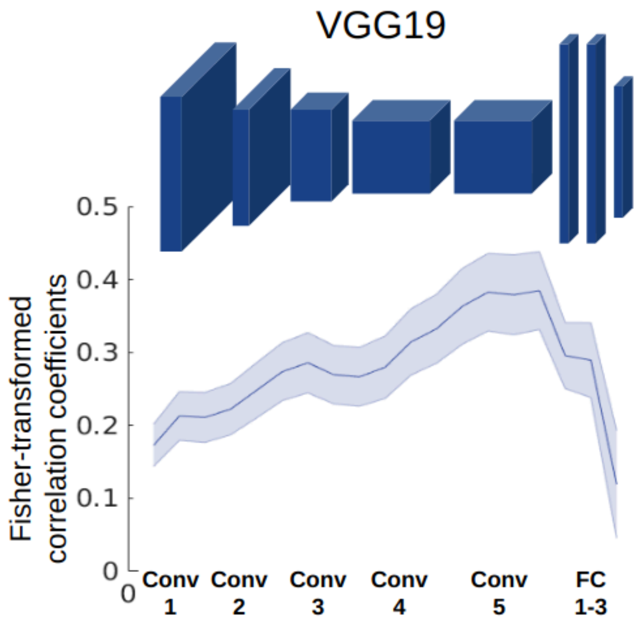


**Figure S3.** (A) Fear profiles of participants can be predicted from the activity generated by the 2700 images in the artificial ('deep') neural network VGG19. By fear profile we mean the different self-reported subjective fear scores over all the animal categories, for an individual participant. Based on the pattern of activity in "latent space"'nodes' within the artificial neural network over many stimuli, we tried to predict these fear profiles for each participant. The r2 coefficientFisher-transformed correlation coefficient is a measure of how well activity from each layer of a network, or activity from the 'latent-space' of thea network (see main text for more details), can accurately predict the fear profile over different animal categories. These results indicate that CLIPboth networks can perform far better than chance (see main text for statistics).

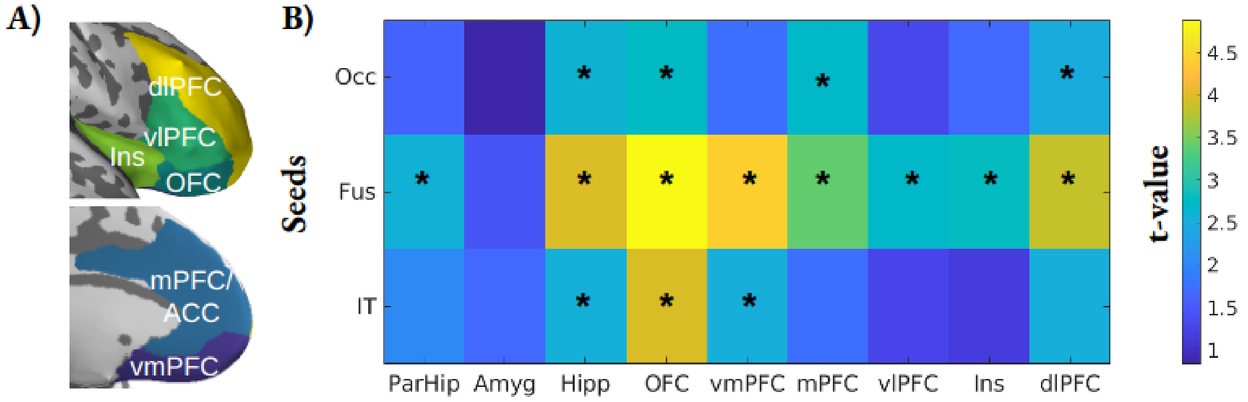


**Figure S4.** Difference in information transmission from ventral visual regions to other brain areas, between participants with and without subjective fear of 'threatening' stimuli. Color coded represent the t-value of the between group difference in a measure of information transmission. Results are processed following the same approach as described in Fig. 4 but using a fixed alpha parameter across ROIs (alpha = 0.1).

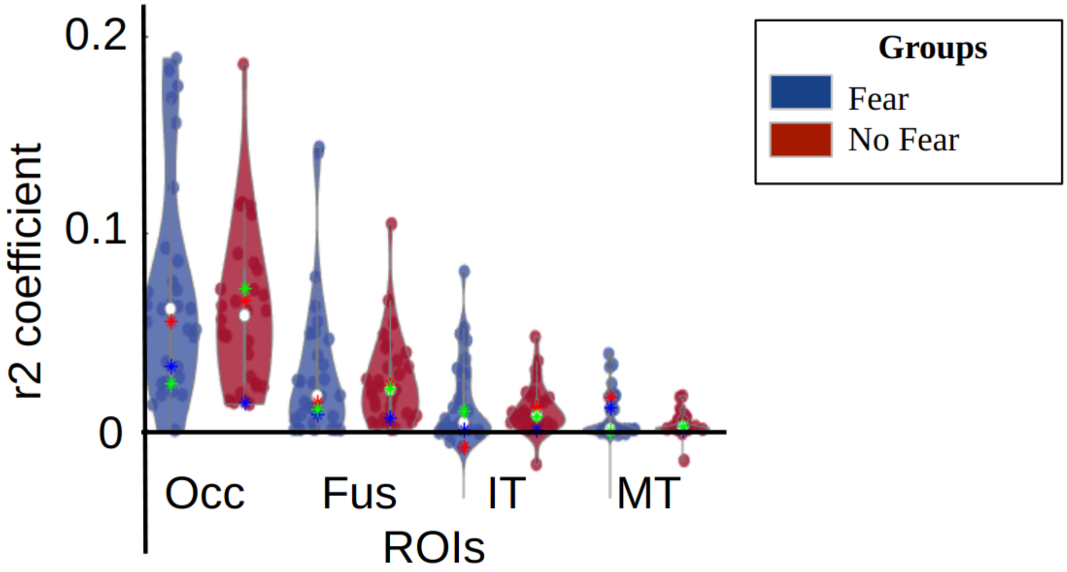


**Figure S5.** Each colored star represents the R2 value of a specific patient included in our sample (N=3). Their z-scores lie between -0.28 and -0.87 in the occipital gyrus and -0.42 and -0.58 in the fusiform gyrus. Overall, they do not represent extreme values in these distributions.

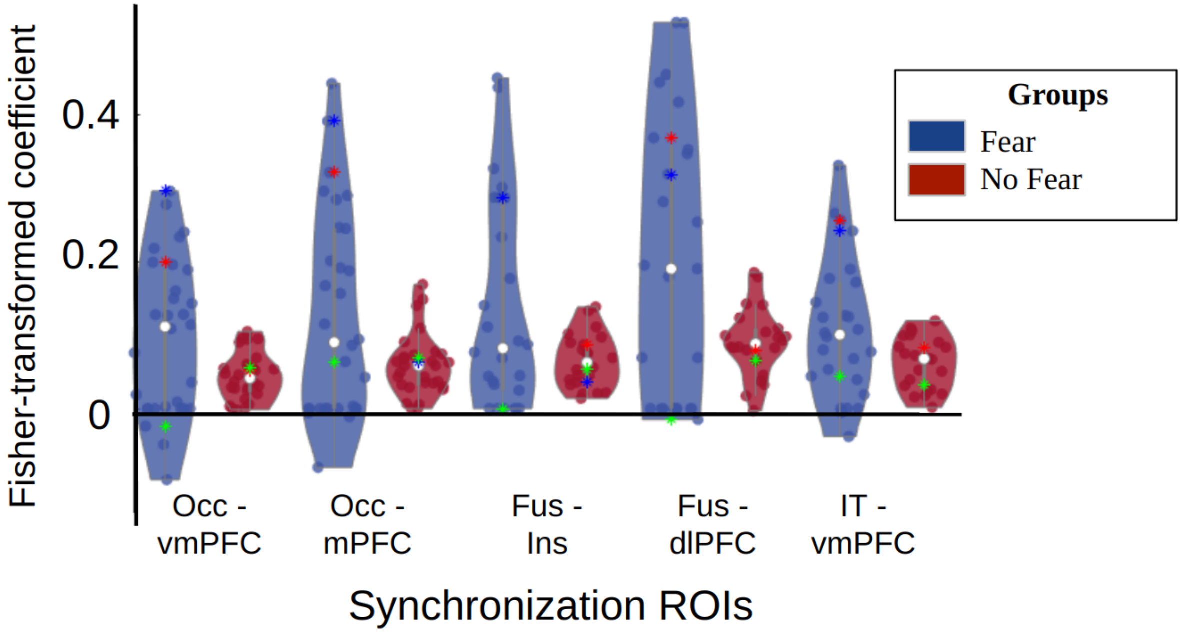


**Figure S6.** Each colored star represents the R2 value of a specific patient included in our sample (N=3). 2 patients (color coded as blue and red stars in the figure) present relatively high synchronization (with multiple instances of synchronization values above 1 STD) in regions of significant synchronization between the ventral visual stream and the prefrontal regions. Paired-sample t-tests remain significant after removing the patients (Occ-vmPFC: $t(25) = 2.91$, $p = 0.0076$; Occ-mPFC: ($t(25) = 2,59$, $p = 0.0159$; Fus-Ins: $t(18) = 2.74$, $p = 0.0135$; Fus-dlPFC: $t(19) = 2.80$, $p = 0.0115$; IT-vmPFC: $t(19) = 2.61$, $p = 0.0173$). However, those results no longer survive a correction for the False discovery rate to correct for the 27 comparisons conducted in the synchronization analyses.
